# *C. elegans* discriminate colors without eyes or opsins

**DOI:** 10.1101/092072

**Authors:** D. Dipon Ghosh, Xin Jin, Michael Nitabach

## Abstract

Here we establish that, contrary to expectations, *Caenorhabditis elegans* nematode worms possess a color discrimination system despite lacking any opsin or other known visible light photoreceptor genes. We found that white light guides *C. elegans* foraging decisions away from harmful bacteria that secrete a blue pigment toxin. Absorption of amber light by this blue pigment toxin alters the color of light sensed by the worm, and thereby triggers an increase in avoidance. By combining narrow-band blue and amber light sources, we demonstrated that detection of the specific blue:amber ratio by the worm guides its foraging decision. These behavioral and psychophysical studies thus establish the existence of a color detection system that is distinct from those of other animals.

## Main text

*C. elegans* live in decomposing organic matter where they feed on microorganisms^1-3^, some of which secrete colorful pigments. While *C. elegans* lack any specialized photoreceptor cells or opsin genes, they possess an illuminance sensing system that mediates rapid escape responses to short-wavelength light^4-6^. However, it is unknown whether *C. elegans* use light information, potentially including color, to inform complex decisions like foraging in environments containing colorful food sources. To address this question, we first tested whether white light alters foraging decisions on *P. aeruginosa* bacterial lawns containing the blue pigment toxin pyocyanin, one of a number of small molecule phenazine toxins secreted by *P. aeruginosa*^7-10^. We employed an 8 kilolux LED array white light source to mimic naturalistic lighting levels (**Fig. 1a, Supplementary Fig. 1a**). Previous studies have shown that foraging decisions to remain on or leave a bacterial lawn are guided by a variety of factors^11, 12^. Worms remain on bacterial lawns that are easy to eat and support growth, while leaving lawns that are of poor nutritive quality, repulsive, or pathogenic^11-16^. Worms are initially attracted to *P. aeruginosa* lawns, but over a time course of hours, as the *P. aeruginosa* continues to divide and secrete toxins, they respond to its increasingly aversive qualities and begin to leave^14, 15, 17-20^. Whether light plays a role in guiding avoidance of *P. aeruginosa* has never been tested. Here we employed a standard lawn avoidance assay, placing worms on a *P. aeruginosa* lawn in the center of an agar plate, and quantified the time course of avoidance as the fraction of worms found off the lawn (**Fig. 1a**). Consistent with prior studies, worms gradually avoid *P. aeruginosa* strain PA14 over a span of many hours^14, 15, 17-20^ (**Fig. 1a**). Surprisingly, however, this avoidance is dramatically potentiated by white light (**Fig. 1b**). White light does not potentiate avoidance of lawns of non-toxic *E. coli* OP50 food bacteria, which worms remain on without leaving (**Fig. 1b**).

**Figure 1.**
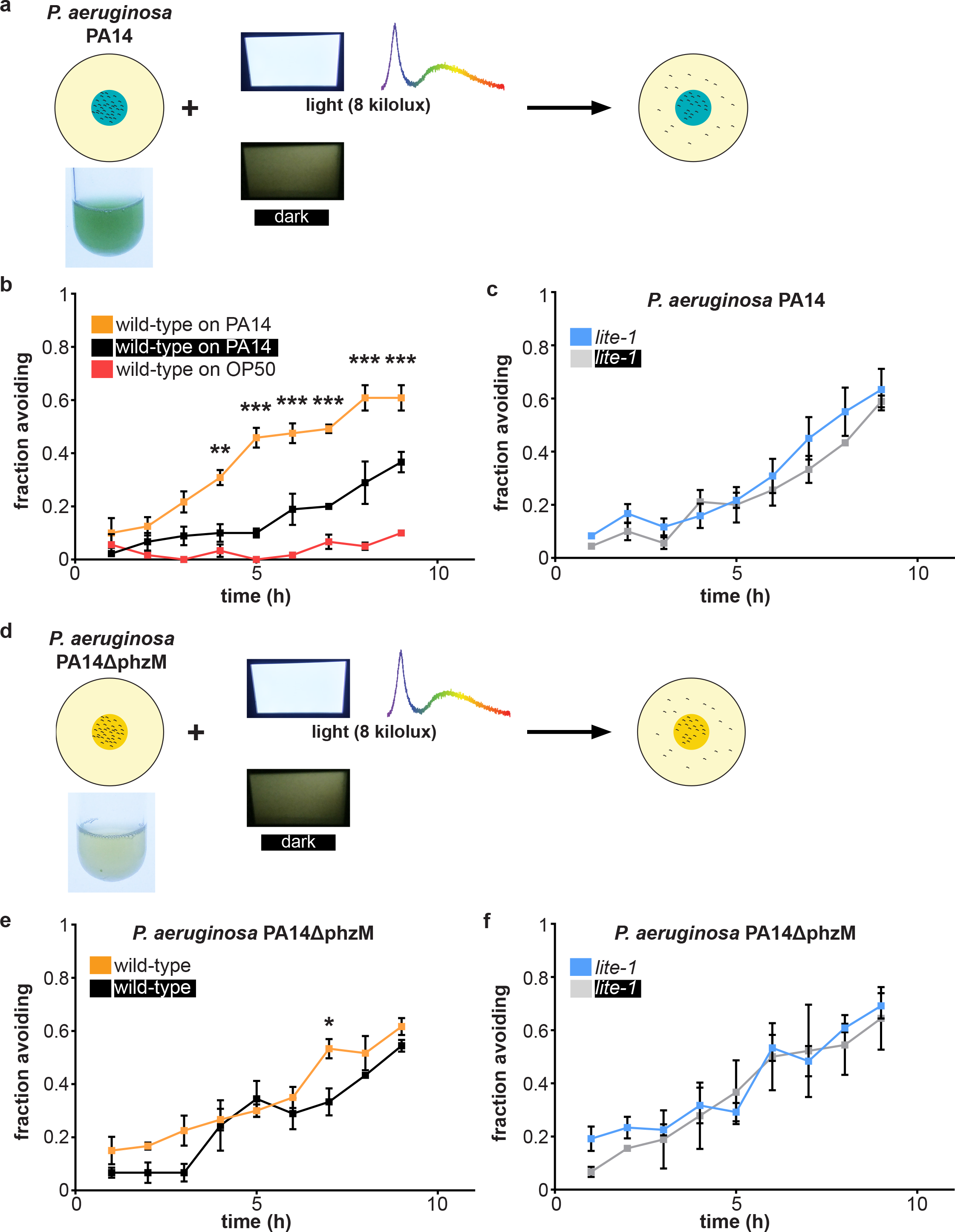
Light-potentiated avoidance of *P. aeruginosa* requires *lite-1* and pyocyanin blue pigment toxin. **(a)** Schematic of bacterial lawn avoidance paradigm. Worms (black squiggles) are placed on a lawn of *P. aeruginosa* strain PA14. Photographs are of *P. aeruginosa* PA14 liquid culture and 8 kilolux white light with corresponding spectrum as measured by a CCD spectrometer. The fraction of worms outside the lawn is counted once per hour in the absence (dark) or presence (light) of 8 kilolux white incident light illuminating the entire plate. Data represent an average of *n* assays with thirty worms per assay. **(b)** Timecourse of wild-type worm avoidance of lawns of PA14 or *E. coli* OP50. Over the course of the nine hour assay, worms increase avoidance of PA14 in the light (orange) and dark (black), but this avoidance is greatly potentiated by white light. Worms remain on OP50 lawns (red) throughout the nine hour assay in the presence of light. *n =* 3-4. **(c)** Avoidance of PA14 by *lite-1* null-mutant worms is unaffected by white light. *n* = 3-4. **(d)** Schematic of lawn avoidance paradigm with *P. aeruginosa* strain PA14ΔphzM, which is incapable of synthesizing pyocyanin. Photographs are of *P. aeruginosa* PA14ΔphzM liquid culture (note the change in color when pyocyanin is absent) and white light used in these experiments. **(e)** White light has a dramatically attenuated effect on avoidance by wild-type worms of PA14ΔphzM. *n =* 3-4. **(f)** Avoidance of PA14ΔphzM by *lite-1* null-mutant worms is unaffected by white light. *n =* 3-4. (All statistical comparisons are by two-way ANOVA with time as a repeated measure and post-hoc Bonferroni tests. For corresponding *p* and F values, see Supplemental Table 1. Error bars denote s.e.m. * indicates *p* < 0.05, ** indicates *p* ≤ 0.01, and *** indicates *p* ≤ 0.001.)

Recent studies have implicated the worm *lite-1* gene – which encodes a non-canonical seven-transmembrane domain protein with closest homology to insect olfactory and gustatory chemoreceptors – in detecting ultraviolet light and mediating photophobic responses to short-wavelength light^4-6, 21, 22^. In addition, *lite-1* is required for short-wavelength light to inhibit feeding through the generation of reactive oxygen species (ROS) like hydrogen peroxide^22^. While the lights used in our experiments do not contain ultraviolet light and purified LITE-1 itself does not absorb visible light^21^, we tested whether the *lite-1* pathway is involved in light-dependent potentiation of PA14 avoidance. Like wild-type worms, *lite-1* null-mutant worms gradually avoid PA14 over time, but their avoidance is unaffected by white light (**Fig. 1c**). These results indicate an essential role for the *lite-1* light response pathway in white light-mediated potentiation of PA14 avoidance.

In order to determine whether the secreted blue toxin pyocyanin is involved in light-dependent potentiation of *P. aeruginosa* avoidance, we tested avoidance of a *phzM* mutant strain, PA14ΔphzM. This strain lacks the biosynthetic enzyme necessary for pyocyanin synthesis, but still synthesizes other phenazine toxins^7-10, 23^, and PA14ΔphzM cultures are no longer blue (**Fig. 1d**). Light has only a minimal effect on avoidance of PA14ΔphzM by wild-type worms, and no effect on avoidance by *lite-1* null-mutant worms (**Fig. 1e, f**). The *phzM* mutation has pleiotropic effects beyond absence of pyocyanin, which include, among other things, increased production and secretion of other phenazines^7^. The results thus far establish light-dependent potentiation of PA14 avoidance requiring both the *lite-1* light response pathway of the worm and the blue toxin pyocyanin secreted by *P. aeruginosa*.

To circumvent complexities associated with dynamic *P. aeruginosa* lawns, which continue to divide and secrete phenazines over the course of the assay, we tested whether pyocyanin is sufficient to mediate light-dependent potentiation of avoidance of otherwise non-toxic bacteria. Accordingly, we employed lawns of non-toxic *E. coli* OP50 supplemented with pyocyanin (**Fig. 2a**). Wild-type worms avoid OP50 supplemented with 2.5 mM pyocyanin within one hour of placing on the lawn, but only in the presence of white light (**Fig. 2b**). This is more rapid than the avoidance of PA14 (see **Fig. 1b**), and is already at the maximum avoidance reached during extended exposure (**Fig. 2b**). This more rapid, unchanging avoidance of OP50 lawns supplemented with 2.5 mM pyocyanin is likely due to the constant amount of pyocyanin present in the lawn, in contrast to *P. aeruginosa* lawns that continue to secrete pyocyanin over the course of the assay. In light of this unchanging avoidance of OP50 lawns supplemented with pyocyanin, for subsequent experiments we relied on this one hour time point. Like avoidance of PA14, light-dependent avoidance of OP50 + 2.5 mM pyocyanin is abolished by the *lite-1* null mutation (**Fig. 2c**). While supplementing with 0.25 mM pyocyanin fails to elicit avoidance, worms avoid OP50 + 5 mM pyocyanin both in the light and dark, and independently of *lite-1* function (**Fig. 2c**). These results indicate that an intermediate concentration of pyocyanin confers light-dependent and *lite-1*-dependent avoidance of otherwise innocuous food bacteria, with higher concentrations conferring avoidance independent of light or *lite-1*.

**Figure 2.**
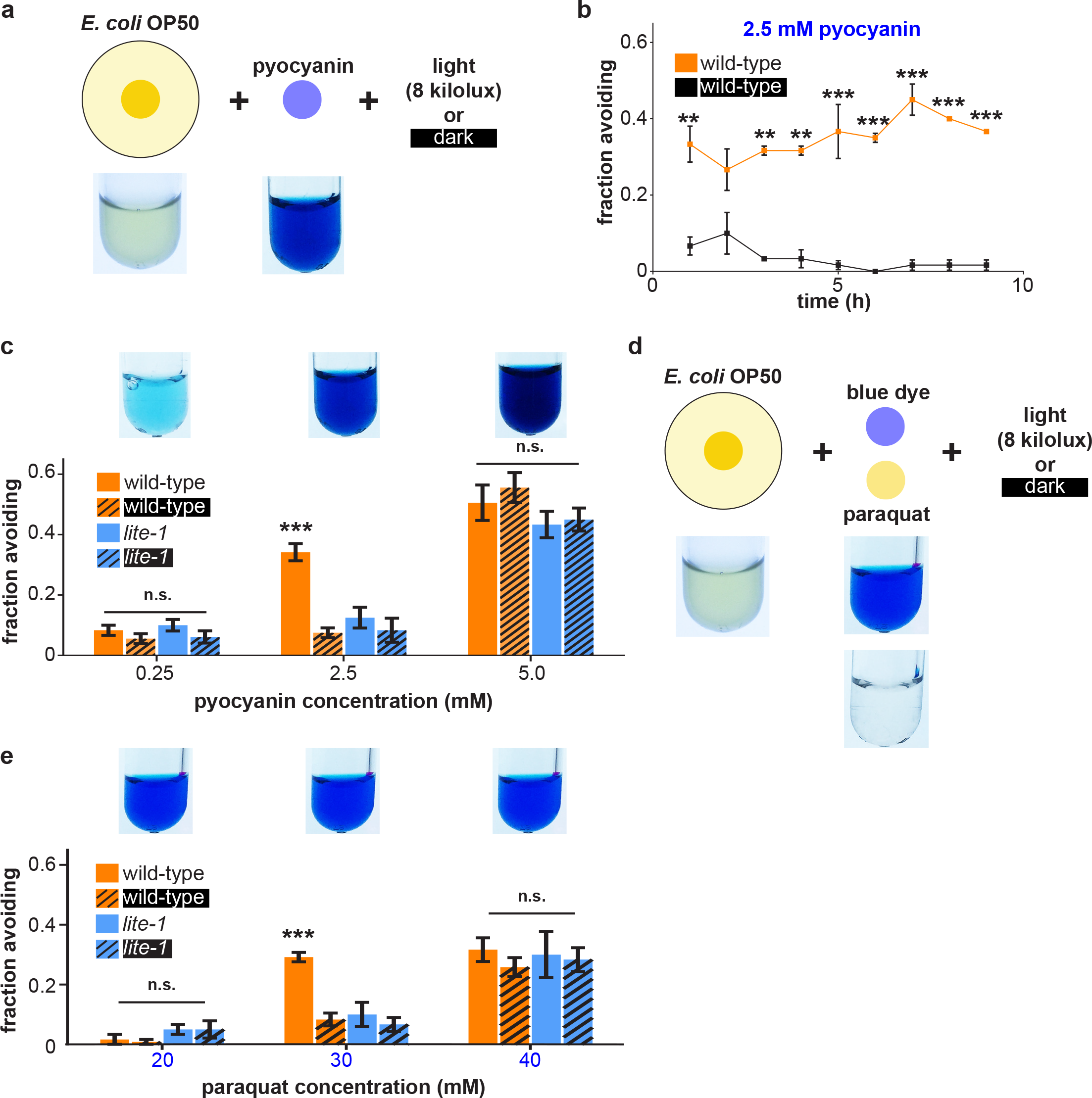
Light potentiates avoidance of otherwise non-toxic *E. coli* lawns containing blue pigment and ROS-generating toxins. **(a)** Schematic of experimental design for testing avoidance of *E. coli* OP50 lawns supplemented with pyocyanin. Photographs are of *E. coli* OP50 liquid culture and test solutions used in these experiments. **(b)** Timecourse of avoidance by wild-type worms of *E. coli* OP50 lawns supplemented with 2.5 mM pyocyanin in the presence or absence of light reveals white light potentiation of stable avoidance. For corresponding *p* and F values, see Supplemental Table 1. *n* = 2-4. **(c)** White light potentiates one-hour avoidance of OP50 lawns supplemented with 2.5 mM pyocyanin by wild-type but not *lite-1* null-mutant worms. Avoidance of OP50 lawns supplemented with 0.25 mM or 5 mM pyocyanin is unaffected by light. *n* = 4. **(d)** Experimental design for testing avoidance of OP50 lawns supplemented with colorless ROS-generating toxin paraquat and inert blue dye. **(e)** White light potentiates avoidance of OP50 lawns supplemented with 30 mM paraquat and blue dye by wild-type but not *lite-1* null-mutant worms. Avoidance of OP50 lawns supplemented with 20 mM or 40 mM paraquat and blue dye is unaffected by light. *n* = 4. (Data in this figure and all remaining figures represent an average of *n* assays with thirty worms per assay. Unless otherwise indicated, all statistical analyses were performed by one-way ANOVA with post-hoc Tukey-Kramer for pairwise comparisons or Dunnett or Bonferroni tests for comparisons with control. Error bars denote s.e.m. * indicates *p* < 0.05, ** indicates *p* ≤ 0.01, and *** indicates *p* ≤ 0.001.)

To test whether the blue color of pyocyanin independent of its toxic chemistry is sufficient to induce avoidance of non-toxic OP50 lawns, we employed OP50 lawns supplemented with non-toxic inert blue dye (**Supplementary Fig. 2a**). Neither wild-type nor *lite-1* null-mutant worms avoid OP50 lawns supplemented with blue dye that visually matches the color of pyocyanin, whether in dark or light, indicating that it is not solely the blue color of pyocyanin that drives light-potentiated avoidance but also its chemical reactivity (**Supplementary Fig. 2b**). Lower and higher concentrations of blue dye also failed to elicit avoidance, whether in dark or light (**Supplementary Fig. 2c**). One of the toxic chemical activities of pyocyanin is that it enters eukaryotic cells and generates reactive oxygen species (ROS)^7-10, 23^. To determine if pyocyanin’s combination of blue color and ROS-generating toxic chemistry underlies light-potentiated avoidance, we employed OP50 lawns supplemented with the colorless ROS-generating toxin paraquat^24^ and non-toxic inert blue dye (**Fig. 2d**). Wild-type worms avoid OP50 supplemented with 30 mM paraquat and blue dye, but only in the presence of white light (**Fig. 2e**). This light-potentiated avoidance is abolished in *lite-1* null-mutant worms (**Fig. 2e**). As with OP50 + pyocyanin, higher concentrations of paraquat mediate avoidance independent of both incident light and *lite-1*, while lower concentrations of paraquat with blue dye are insufficient to mediate avoidance (**Fig. 2e**). Supplementing OP50 lawns with 30 mM paraquat without any dye or with inert non-toxic red dye (**Supplementary Fig. 2d**) are each insufficient to confer light-potentiated avoidance (**Supplementary Fig. 2e**). These results indicate that light- and *lite-1*-dependent avoidance of pyocyanin-containing bacterial lawns relies both on its chemical ROS-generating reactivity and on its blue color. They also indicate that avoidance conferred by higher concentrations of pyocyanin relies solely on its ROS-generating reactivity and is independent of *lite-1* function. A recent study determined that the *gur-3* gene – which encodes another *Drosophila* gustatory receptor paralog related to *lite-1* – mediates the detection and response to ROS, including hydrogen peroxide^22^. We therefore tested *gur-3* null-mutant worms for light-potentiated avoidance of OP50 lawns supplemented with paraquat and blue dye. Surprisingly, *gur-3* null-mutant worms avoid lawns supplemented with paraquat and blue dye more than wild-type worms in the dark, and this enhanced avoidance is suppressed by light (**Supplementary Fig. 2f**). This potentiated avoidance in the dark is suppressed in *lite-1; gur-3* double mutant worms (**Supplementary Fig. 2f**). These results reveal a contribution of *gur-3* to light-potentiated avoidance of ROS-generating toxins, and indicate unresolved complexity of underlying genetic architecture.

How does the presence of blue pigment confer light-potentiated avoidance of bacterial lawns containing ROS-generating toxins? We hypothesized it is by absorbing long-wavelength light and thereby altering the spectral composition of light sensed by the worm. Indeed, white light filtered through either pyocyanin or blue dye solutions exhibits a reduction in long-wavelength content (**Supplementary Fig. 3**). To test this hypothesis, we employed a series of shortpass and longpass optical filters to alter the spectrum of incident light (corresponding photographs and spectra of filtered light are shown in **Supplementary Fig. 4a, b**). Not surprisingly, in light of the previously determined wavelength dependence of *lite-1*-mediated photophobic responses^4-6^, longpass-filtered light lacking blue photons fails to potentiate avoidance of OP50 + 2.5 mM pyocyanin (**Fig. 3a**). Surprisingly, however, shortpass-filtered light with cut-offs of <500 nm or <550 nm also failed to potentiate avoidance (**Fig. 3a**). Only shortpass-filtered light with <600 nm cut-off, which still includes the yellow-orange (amber) component of white light, was sufficient to potentiate avoidance of OP50 + 2.5 mM pyocyanin (**Fig. 3a**). This establishes that light-potentiated avoidance of bacterial lawns containing ROS-generating toxins and blue pigment requires not only short-wavelength blue light, but also long-wavelength amber light. This suggests the existence of an amber sensing pathway in addition to a blue sensing pathway.

**Figure 3.**
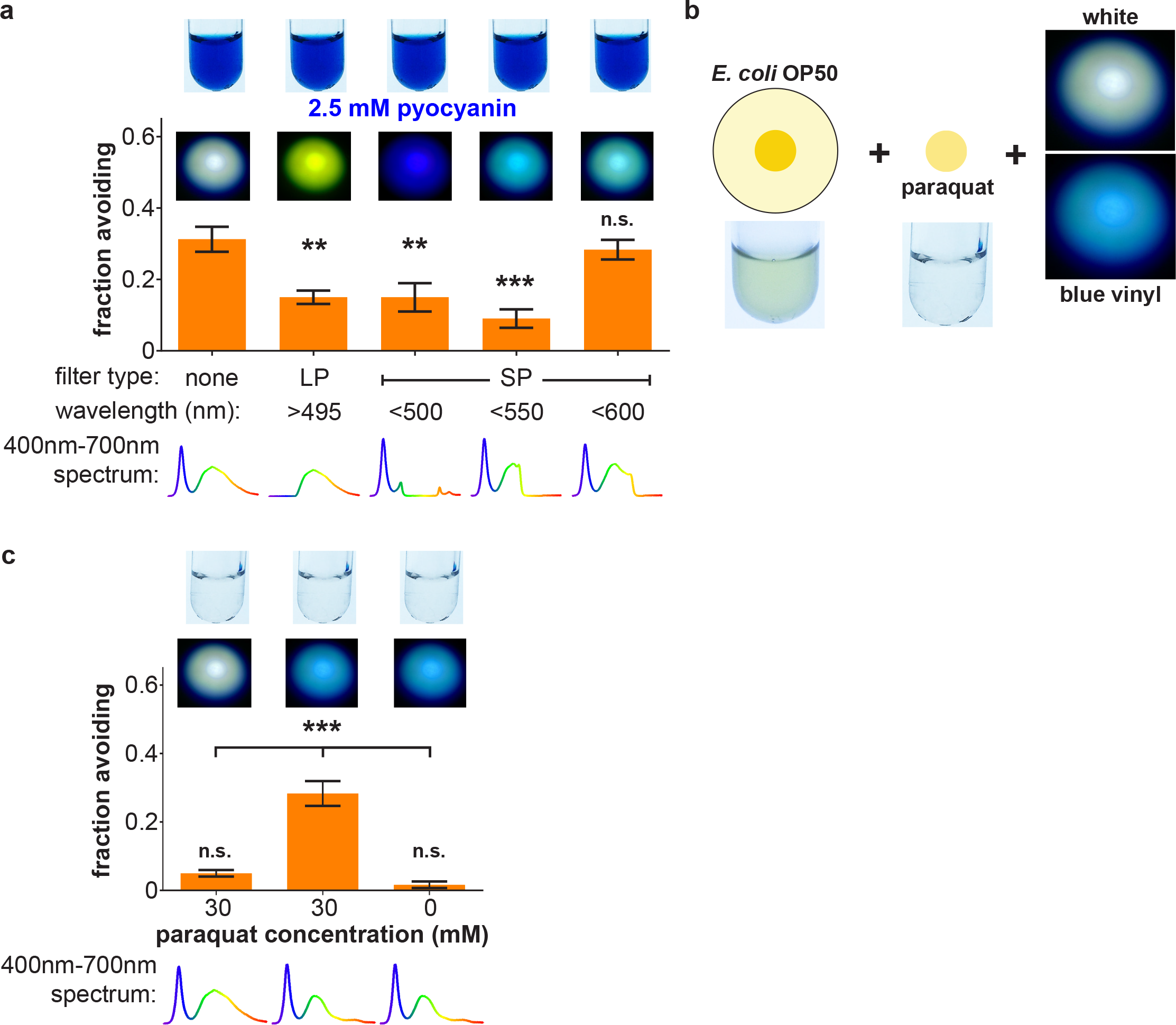
Color of light determines avoidance of toxic bacterial lawns. **(a)** The spectral composition of light was modified by various optical filters. Photographs are of *E. coli* OP50 liquid culture and test solutions used in these experiments, as well as lights modified by the indicated filter shining on a white background. Light spectra were measured using a CCD spectrometer. Blue light or blue-green light obtained with 500 nm or 550 nm shortpass filters and yellow-green light obtained with a 495 nm longpass filter each fail to potentiate avoidance of OP50 lawns supplemented with 2.5 mM pyocyanin. Only light containing both blue and amber light, obtained with a 600 nm cut-off shortpass filter, potentiates avoidance of OP50 supplemented with 2.5 mM pyocyanin. *n* = 4-8. **(b, c)** Light modified by a blue vinyl filter that decreases intensity of amber light without eliminating it potentiates avoidance of colorless OP50 lawns supplemented with 30 mM paraquat, but not lawns lacking paraquat. *n* = 4-6.

Completely eliminating amber light abolishes light-dependent potentiation of avoidance of OP50 + 2.5 mM pyocyanin (**Fig. 3a**). Thus, we hypothesized that it is the decreased ratio of amber to blue light caused by the blue pigment (see **Supplementary Fig. 3**) that potentiates avoidance of bacterial lawns containing ROS-generating toxins. To test this hypothesis, we eliminated blue pigment from the lawn and directly modified the spectral composition of light sensed by the worm with a “blue vinyl” filter selected to match the absorbance of blue pigment that reduces without eliminating amber photons (**Fig. 3b, Supplementary Fig. 4c**). Indeed, light filtered through the blue vinyl filter recapitulates white light-potentiated avoidance of OP50 + 30 mM paraquat, but now in the absence of blue pigment in the lawn (**Fig. 3c**). These results establish that blue pigments confer light-potentiated avoidance of toxic bacterial lawns by absorbing amber light and thereby altering the spectral composition of light sensed by the worm.

**Figure 4.**
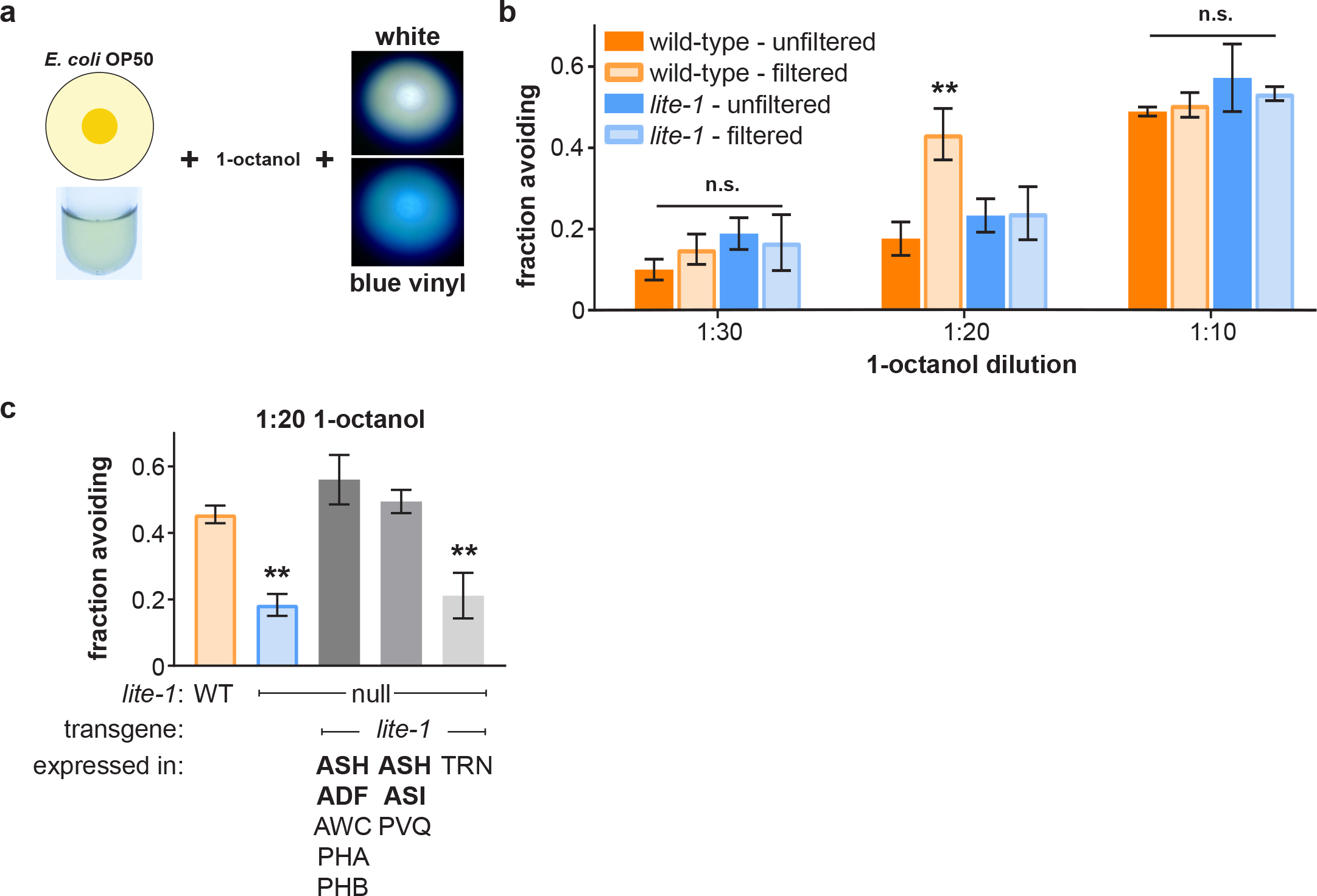
Color of light determines bacterial lawn avoidance in the presence of the aversive odorant 1-octanol. **(a, b)** Blue vinyl-filtered light potentiates avoidance of OP50 lawns in the presence of 1:20 1-octanol by wild-type but not *lite-1* null-mutant worms. Avoidance of OP50 lawns in the presence of 1:30 or 1:10 1-octanol is unaffected. *n* = 6. **(c)** Re-expression of *lite-1* in *lite-1* null-mutant worms using the *gpa-13* or *sra-6* promoters for expression in neurons ASH/ADF/AWC/PHA/PHB and ASH/ASI/PVQ, respectively, rescues color-dependent avoidance of OP50 lawns. Re-expression of *lite-1* using the *mec-17* promoter for expression in touch receptor neurons (TRN) has no effect. *n* = 6.

To further confirm that it is the altered color of light, specifically the blue-to-amber ratio, that potentiates avoidance and not altered illuminance, we varied the intensity of simulated daylight (**Supplementary Fig. 5a**). Increasing white light intensity by ∼3-fold to 25 klx failed to increase avoidance of OP50 + 30 mM paraquat (**Supplementary Fig. 5b**). While further increases to 50 klx or 100 klx did mediate increased avoidance of OP50 + 30 mM paraquat, these very high light intensities closer to that of direct sunlight (100-120 kilolux) identically increase avoidance of OP50 lawns without paraquat, most likely by triggering *lite-*1-dependent escape responses (**Supplementary Fig. 5b**). Since *lite-1*-dependent responses to light may themselves involve ROS^22, 25^, we used a hydrogen peroxide sensor to determine if 8 klx light induces ROS production. No hydrogen peroxide is detected when worms, OP50 bacteria, paraquat, and blue dye were combined in a test tube and illuminated with 8 klx light (**Supplementary Fig. 5c**). In contrast, a recent study demonstrated that brighter light (13 mW/mm^2^ of 436 nm blue light) generates approximately 50 µM hydrogen peroxide in a solution containing riboflavin^22^. In addition, this study also demonstrated that at least 10 µM applied hydrogen peroxide is necessary to inhibit worm feeding^22^. We also used H2DCFDA, a cell-permeant fluorescent ROS indicator, to determine if intensities of light used in our experiments induce detectable ROS production in worms. No ROS signal is detected in worms placed on OP50 lawns supplemented with 0 or 30 mM paraquat in the dark or exposed to 8 klx white light for one hour (**Supplementary Fig. 5d, e**). In contrast, substantial ROS signal is detected in worms on OP50 lawns exposed to 100 klx white light for one hour, regardless of whether paraquat is present (**Supplementary Fig. 5d, e**). These results rule out the possibility that blue pigment potentiates avoidance of bacterial lawns containing ROS-generating toxins by scattering blue light and thereby increasing short-wavelength photon flux through the worm. We also tested the effect of light and color in the absence of food bacteria. Neither the presence of light nor changing the color of light affects chemotaxis to the attractive odorant diacetyl or avoidance of an aversive hyperosmotic fructose barrier (**Supplementary Fig. 6a, b**). When given a choice (**Supplementary Fig. 6c**), worms prefer blue vinyl-filtered light over unfiltered white light (**Supplementary Fig. 6d**), although this could be attributable to a preference for dimmer light. Taken together, these results strongly support the conclusion that color-dependent potentiation of avoidance of toxic bacterial lawns operates by a different mechanism than the photophobic escape responses to light.

**Figure 5.**
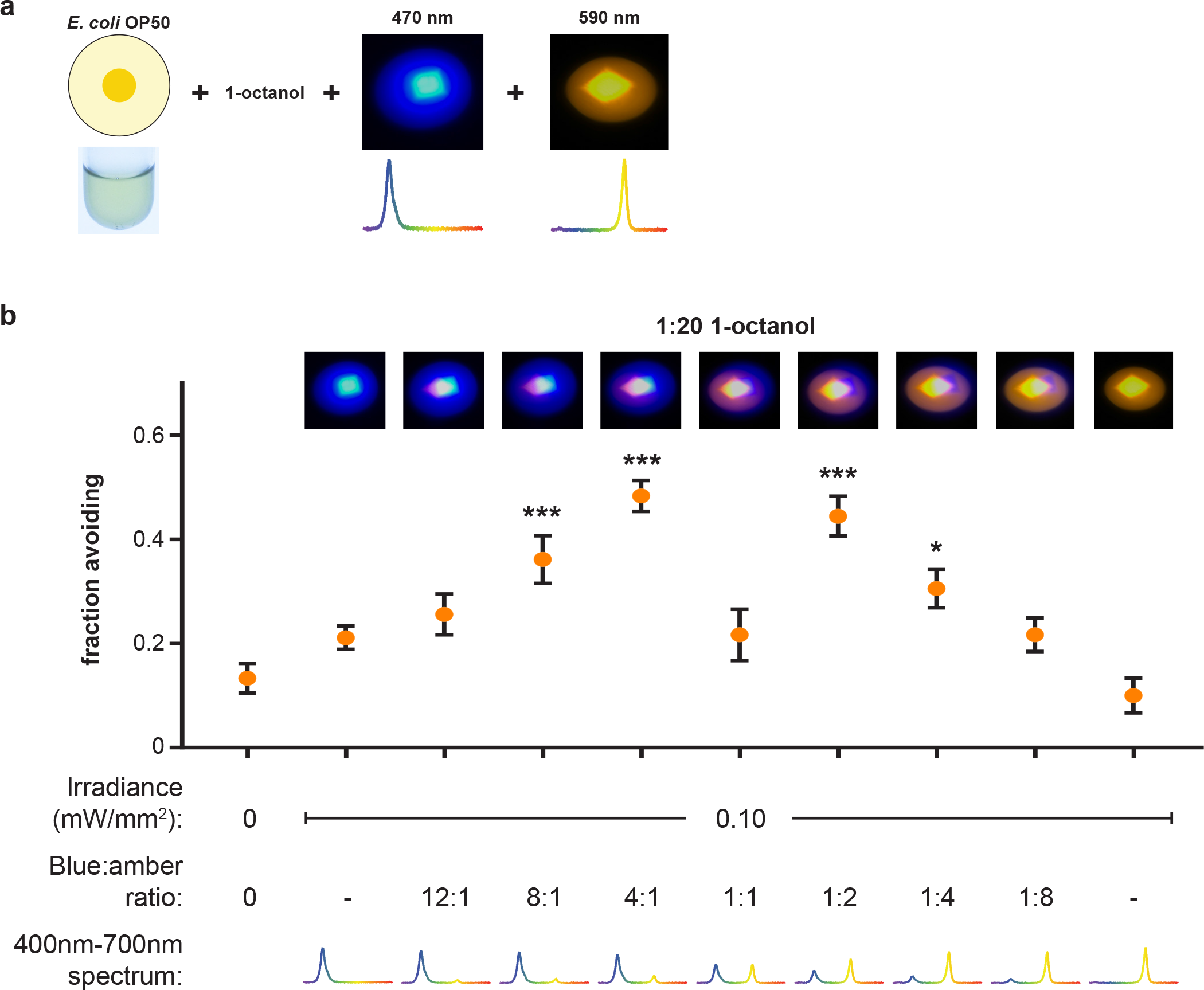
Ratio of narrow-band blue and amber light intensities determines potentiation of lawn avoidance. **(a, b)** The spectral composition of light (blue:amber ratio) was controlled by varying intensity of two narrow-band light sources, keeping total intensity constant at 0.10 mW/mm^2^. Blue light, amber light, or a 1:1 blue:amber fail to potentiate avoidance of OP50 lawns in the presence of 1:20 1-octanol. However, 4:1 or 1:2 blue:amber light potentiate avoidance, while increasing blue:amber beyond 4:1 or decreasing blue:amber below 1:2 attenuates and ultimately eliminates potentiation of avoidance. *n* = 6.

To determine whether color-dependent potentiation of lawn avoidance is specific to the context of ROS-generating toxins, we tested for color-dependent potentiation of avoidance of OP50 lawns in the context of the aversive odorant 1-octanol^26, 27^, a tiny droplet (1 µL) of which was applied to a glass lid covering the assay plate (**Fig. 4a**). Similar to OP50 lawns supplemented with ROS-generating toxins, blue vinyl-filtered light potentiates wild-type worm avoidance of OP50 lawns in the presence of 1:20 octanol compared to unfiltered white light, but not lower or higher dilutions (**Fig. 4b**). This color-dependent lawn avoidance requires *lite-1* (**Fig. 4b**), but interestingly not *gur-3* (**Supplementary Fig. 7**), indicating that the color-dependent responses to ROS-generating toxins (**Supplementary Fig. 2f**) and non-ROS-generating aversive stimuli rely on distinct genetic pathways, although both include *lite-1*. These results demonstrate that color-dependent potentiation of lawn avoidance is not specific to ROS-generating toxins, does not involve previously demonstrated ROS-induced alterations on ASH-mediated sensory responses^28, 29^, and generalizes to other aversive stimuli that do not operate through generation of ROS.

*lite-1* is expressed in a number of neurons in the worm nervous system, including several primary sensory neurons: ASJ, ASH, and ASK^5, 6, 22^. ASJ has been previously implicated in avoidance of PA14 mediated by direct sensory responses to phenazines that induce Ca^2+^ increases and gene activation^15^. In addition, *lite-1* expression in ASJ and ASK is sufficient for escape responses to short-wavelength light^5, 6^. To determine the cellular locus of *lite-1* expression responsible for light-dependent avoidance of OP50 + 2.5 mM pyocyanin, we employed *lite-1* null-mutant worms with targeted re-expression of *lite-1* in neurons ASJ, ASH/ASI/PVQ, or ASK. Restoration of *lite-1* expression in ASJ or ASH/ASI/PVQ are each sufficient to rescue light-dependent avoidance of OP50 + 2.5 mM pyocyanin, while restoration in ASK is not (**Supplementary Fig. 8a**). ASH sensory neurons mediate acute avoidance of octanol^26, 27^, so we also tested whether *lite-1* re-expression in ASH in *lite-1* null-mutant worms is sufficient to rescue color-dependent lawn avoidance in the presence of octanol. Re-expression of *lite-1* using either of two cell-specific promoters whose expression overlaps only in ASH rescues color-dependent lawn avoidance in the presence of octanol (**Fig. 4c**). Re-expression using a promoter active in touch receptor neurons, which do not normally express *lite-1*, has no effect (**Fig. 4c**). While these results do not rule out the contribution of other *lite-1*-expressing neurons, together they indicate that ASJ and ASH are independently sufficient cellular loci important for color discrimination behaviors.

To unambiguously and definitively establish that worms possess two distinct light sensing channels centered at blue and amber wavelengths, respectively, we employed narrowband light sources with peaks at 470 nm (blue) and 590 nm (amber). We combined these lights in different intensity ratios, but with total intensity held constant at 0.10 mW/mm^2^, and tested for potentiation of OP50 lawn avoidance in the presence of 1:20 octanol (**Fig. 5a**). Pure blue light, pure amber light, and 1:1 blue:amber light each fail to potentiate lawn avoidance (**Fig. 5b**). In contrast, 4:1 blue:amber light potentiates avoidance to a similar extent as blue vinyl-filtered white light (**Fig. 5b**). Potentiation of avoidance attenuates as blue:amber ratio increase to 8:1 and is eliminated at 12:1 (**Fig. 5b**). These results are consistent with potentiation of lawn avoidance by blue vinyl-filtered white light, which also increases the blue:amber ratio (see **Supplementary Fig. 3, 4**). Remarkably, decreasing the blue:amber ratio to 1:2, so that amber light now predominates, also potentiates avoidance (**Fig. 5b**). Further decreases to 1:4 and 1:8 progressively attenuate and then eliminate this potentiation (**Fig. 5b**). These psychophysical studies demonstrate that worms can compare the activation of blue- and amber-sensitive light detection channels to potentiate lawn avoidance when the ratio of activation deviates in either direction from a point of equipoise. This establishes that *C. elegans*, despite lacking opsins or other visible-light photoreceptor proteins, are capable of color discrimination.

Taken together, our studies reveal the unexpected influence of color on a *C. elegans* foraging decision. Based on our genetic, behavioral, and psychophysical studies, we speculate that the integration of color and chemical information could enable more accurate discrimination of toxic from non-toxic lawns during foraging by increasing sensitivity and thereby responsiveness to noxious stimuli in the immediate environment (**Supplementary Fig. 9**). If so, color detection potentially provides another input into multisensory integration processes that can refine natural *C. elegans* behaviors^30-34^. Future studies are required to address the cellular and molecular mechanisms underlying the blue and amber light-sensing channels and their integration to guide decision making, as well as to determine whether this color discrimination system might be used to detect other features of the environment, such as sky color as a cue of time of day^35^.

## Acknowledgments

We thank S. Xu and the Caenorhabditis Genetics Center for *C. elegans* strains, F. Ausubel for *P. aeruginosa* strains, and members of the Nitabach and Kazmierczak labs and the Yale worm community for technical support, advice, and comments. D.D.G was supported by a National Institute of Neurological Disease and Stroke (NINDS) National Institutes of Health (NIH) Predoctoral Fellowship (F31NS080628). Work in the laboratory of M.N.N. was supported in part by NINDS, NIH (R01NS055035, R01NS056443, R01NS091070) and National Institute of General Medical Sciences, NIH (R01GM098931).

## Author contributions

D.D.G and M.N.N designed the study and experiments. D.D.G and X.J. performed experiments. D.D.G, X.J., and M.N.N analyzed the data. D.D.G and M.N.N wrote the paper.

## Supplementary Materials

Methods

Supplementary Table 1

Supplementary Figures 1-9

References (including new reference *36*)

